# Stability of host-parasite systems: you must differ to coevolve

**DOI:** 10.1101/400150

**Authors:** Faina Berezovskaya, Georgy P. Karev, Mikhail I. Katsnelson, Yuri I. Wolf, Eugene V. Koonin

## Abstract

**Background:** Genetic parasites are ubiquitous satellites of cellular life forms most of which host a variety of mobile genetic elements including transposons, plasmids and viruses. Theoretical considerations and computer simulations suggest that emergence of genetic parasites is intrinsic to evolving replicator systems.

**Results:** Using methods of bifurcation analysis, we investigated the stability of simple models of replicator-parasite coevolution in a well-mixed environment. It is shown that the simplest imaginable system of this type, one in which the parasite evolves during the replication of the host genome through a minimal mutation that renders the genome of the emerging parasite incapable of producing the replicase but able to recognize and recruit it for its own replication, is unstable. In this model, there are only either trivial or “semi-trivial”, parasite-free equilibria: an inefficient parasite is outcompeted by the host and dies off whereas an efficient one pushes the host out of existence, which leads to the collapse of the entire system. We show that stable host-parasite coevolution (a non-trivial equilibrium) is possible in a modified model where the parasite is qualitatively distinct from the host replicator in that the replication of the parasite depends solely on the availability of the host but not on the carrying capacity of the environment.

**Conclusions:** We analytically determine the conditions for stable host-parasite coevolution in simple mathematical models and find that a parasite that initially evolves from the host through the loss of the ability to replicate autonomously must be substantially derived for a stable host-parasite coevolution regime to be established.

## Background

Genetic parasites are ubiquitous among cellular organisms [1]. In fact, most organisms host a variety of mobile genetic elements (MGE) that differ in their reproduction strategies and the mode of parasite-host interaction, including transposons, plasmids and viruses [2]. The abundance of the MGE in the biosphere is enormous. Viruses are by far the most common biological entities on earth [3-6], genes of MGE, such as those encoding transposases, are among the most abundant ones in diverse environments [7-9], and the genomes of many multicellular organisms consist of up to 50% integrated MGE, in the case of mammals, or even up to 90% in the case of plants [10, 11].

The entire history of life can be properly depicted only as the perennial coevolution of cellular organisms with genetic parasites that includes both the proverbial arms race and various forms of cooperation [1, 12, 13]. Moreover, multiple lines of evidence point to a major role of genetic parasites in the evolution of biological complexity, in general, and in major transitions in evolution, in particular [14, 15].

The ubiquity and the enormous abundance of the MGE in the biosphere imply that genetic parasitism is an intrinsic feature of life. Indeed, parasites invariably emerge in computer simulations of the evolution of simple replicator systems which, in well-mixed models, typically leads to the collapse of the entire system [16-20]. This outcome can be avoided by incorporating compartmentalization into the model [18-20]. Further, mathematical models of the evolution of genomes with integrated MGE, combined with probabilistic reconstruction of the evolution of bacterial and archaeal genomes, suggest that horizontal gene transfer at rates that are required to stave off the mutational meltdown of microbial populations (Muller’s ratchet) prevents elimination of genetic parasites [21].

Cellular organisms and genetic parasites have been considered the two “empires” of life that fundamentally differ with regard to the capability of autonomous reproduction [22, 23]. The MGE fully depend on the host for energy production and the biosynthetic processes, in particular, translation (notwithstanding the fact that many large viruses encode components of the respective functional systems that modify and modulate the respective host functions [24, 25]).

In our previous theoretical analysis of the evolution of genetic parasites in simple replicator systems [26] which was, to a large extent, inspired by the seminal early experiments of Spiegelman and colleagues on reductive evolution of bacteriophages genomes *in vitro* [27-30], we present a semi-formal argument that the parasite-free state of a replicator is inherently evolutionarily unstable. Here, we explore these and derivative models analytically and show that, in order for a replicator and a parasite to stably coevolve, the parasite must substantially differ from the host in its reproduction strategy.

## Results and Discussion

### Genetic parasite as a degraded variant of the replicator

The simplest conceptual model of the emergence of a genetic parasite from within a self-replicating system [26] implies (initially infinitesimal) degradation of the replicase-encoding signal while the replicase-recognition signal is retained. The dynamics of such a system is described by the following pair of ordinary differential equations [26]:

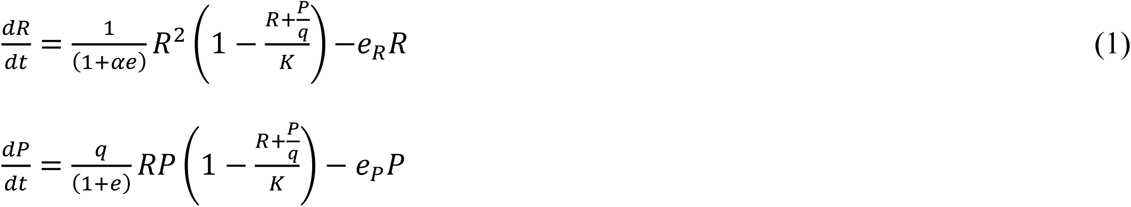

(see Table 1 for the model parameters). The model is based on the following assumptions:

• The kinetics of replication depends on the interaction of the replicator and the template (*R*^2^ for the replicator and *RP* for the parasite)

• The decay rates are constant (*e*_*r*_ and *e*_*P*_) for replicator and parasite respectively

• The parasite template replicates faster than that of the replicator by the factor *q* ≥ 1, which, as the first approximation, could interpreted as the “economy” factor (the parasite being *q* times smaller than the replicator)

• Both populations are environmentally limited by the same resources; with the parasites more economical by the same factor *q*

• The replicators possess a (costly) defense mechanism with efficiency *e* ≥ 0 that is capable to suppress the parasite replication by a factor of 1 + *e* at the cost to the replicators replication rate of 1 + *αe* (where *α* ≥ 0 is the defense mechanism cost factor)

As noticed previously [26], this system is unstable at the point (*R* ≅ *K*, *P* = 0, *e* = 0), that is, introduction of the parasite into a replicator system without any defense near the equilibrium leads to runaway parasite proliferation and the eventual collapse of the entire system if *q* > *e*_*P*_/*e*_*R*_.

Here, we aim to identify all stationary states of model (1) and to analyze their stability, to study the qualitative behavior of this model and to suggest some modifications that make possible the stable host-parasite co-evolution.

All possible equilibria of the model can be found from the system of equations 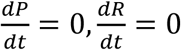. These equations determine the following pair of (non-zero) isoclines (see Additional File 1 for details):

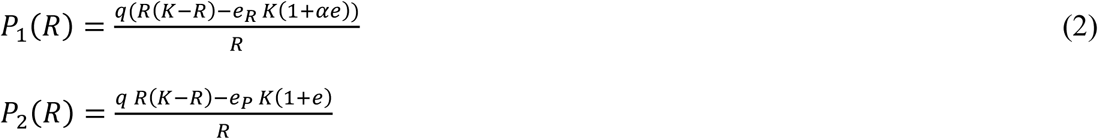

the intersection of which (*P*_1_(*R*) = *P*_2_(*R*) at *R* > 0, *P* > 0) indicates a non-trivial equilibrium. It should be immediately apparent, however, that both equations have the same form and differ only by the values of the coefficients *q*(1 + *αe*)*e*_*R*_ and (1 + *e*)*e*_*P*_, respectively (see Figure 1, left panel, for the characteristic shapes of these isoclines when *q*(1 + *αe*)*e*_*R*_ ≠ (1 + *e*)*e*_*P*_). Therefore, the curves do not intersect in any point (*R* > 0, *P* > 0); the only case where such equilibria exist is 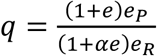 ; in this case *P*_1_(*R*) = *P*_2_(*R*) over the whole range of *R* and the system has a line of non-isolated equilibria (see Figure 1, right panel).

**Figure 1.**
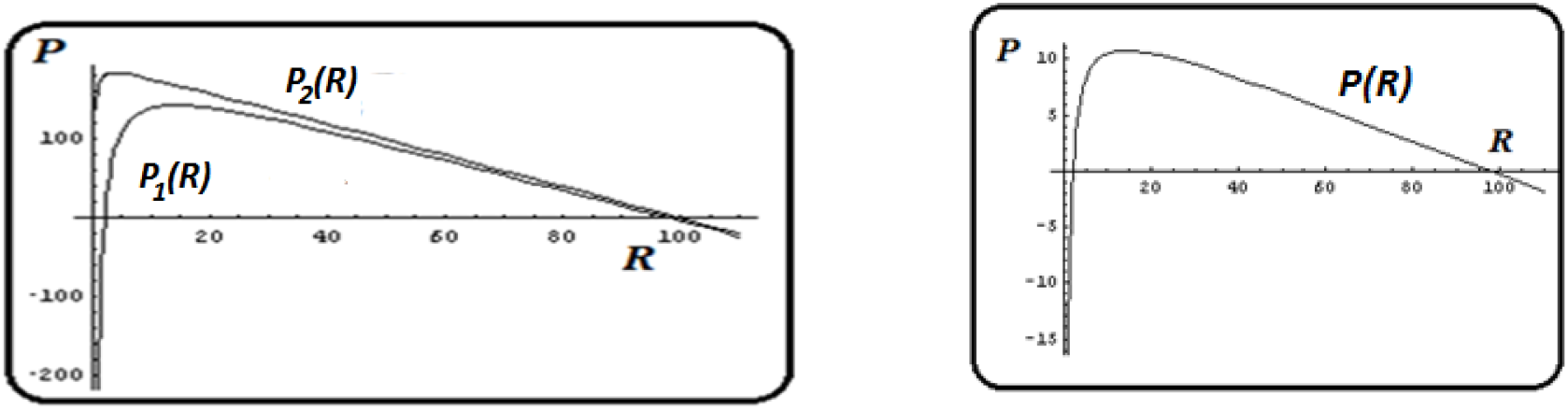
Null isoclines for model (1). Left panel: isoclines (equation (2); right panel: curve of non-isolated equilibria.

More formally, model (1) always has the trivial equilibrium *O* (*R* = 0, *P* = 0) (both populations collapse). In addition to that, the model might have up to two parasite-free, “semi-trivial” equilibria

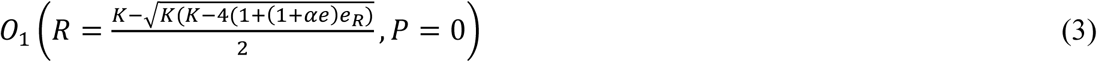

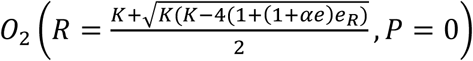

the existence of which depends on the value of the following expression:

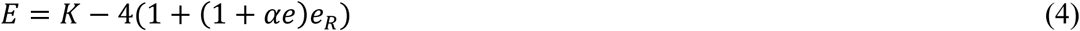

If *E* > 0, both *O*_1_ and *O*_2_ exist; if *E* = 0, *O*_1_ = *O*_2_ = *O*_12_(*R* = *K*/2, *P* = 0); if *E* < 0, the model has only the trivial equilibrium *O*.

Stability analysis shows that the trivial equilibrium *O* is a stable node at all values of the model parameters. When both *O*_1_ and *O*_2_ exist (*E* > 0), if 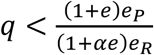, *O*_1_ is a saddle and *O*_2_ is a stable node and if 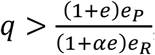, *O*_1_ is an unstable node and *O*_2_ is a saddle. If *k* = 4(1 + *e*)/*q*, the only semi-trivial equilibrium *O*_12_ is a saddle-node fixed point (Figure 2b; see Additional File 1 for details). Let us consider *q* and *e* as parameters of model (1) whereas *k*, *α*, *e*_*R*_, *e*_*P*_ are arbitrary values of fixed coefficients. There exist two qualitatively different types of asymptotical (*t* → ∞) behavior of the system, namely:

**Figure 2.**
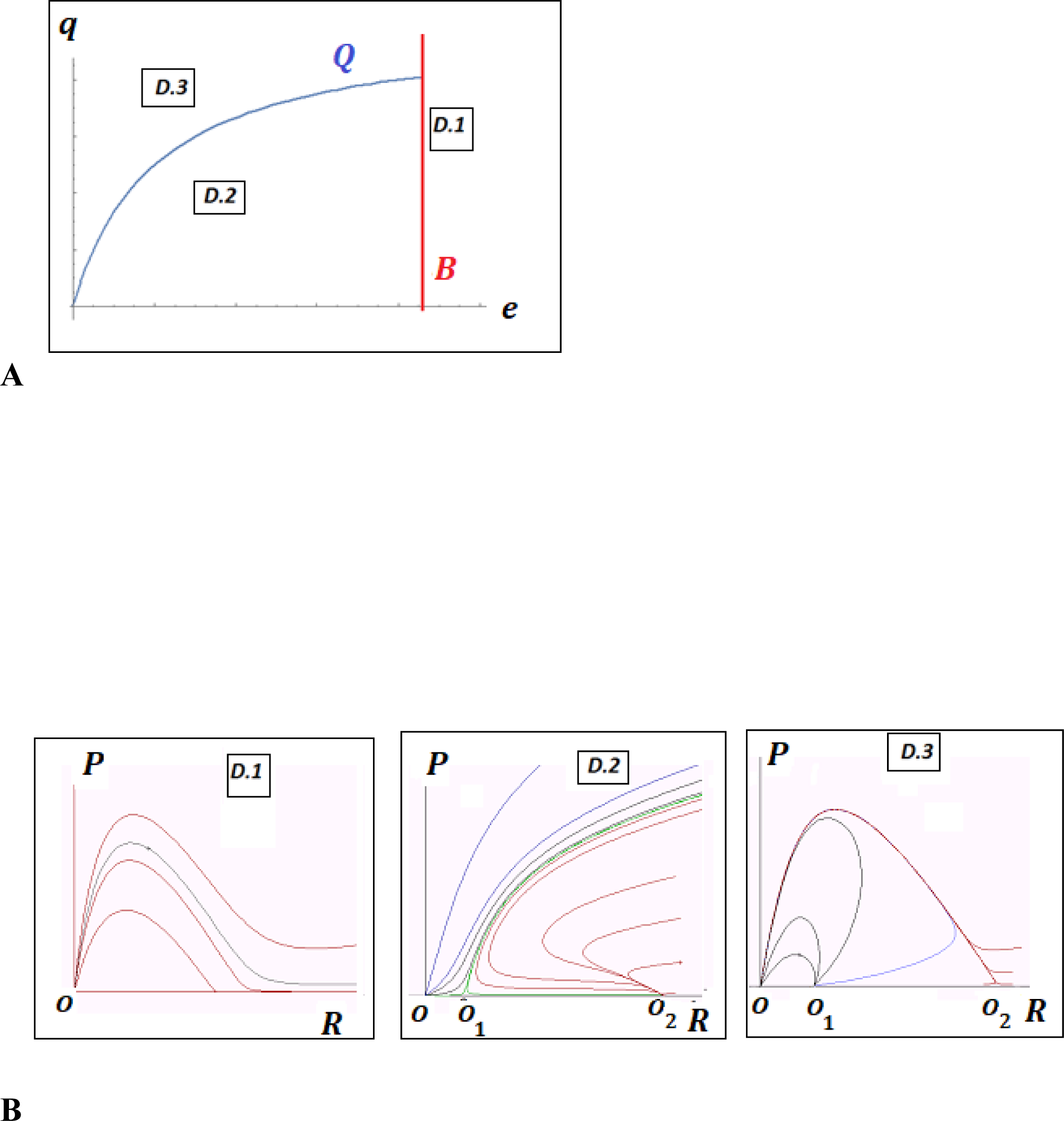
Characteristics of model (1). **A** Parameter-phase portrait. ***D*_1_**, ***D*_2_** and ***D*_3_** indicate domains with qualitatively different model behaviors. The boundaries are: ***Q***: 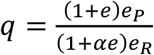(blue), ***B***: 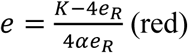. **B** Examples of phase portraits of model (1) with a stable trivial equilibrium *O* (all domains), unstable semi-trivial equilibrium *O*_1_ (domains ***D*_2_** and ***D*_3_)** and stable semi-trivial equilibrium *O*_2_ (domain ***D*_2_**).

1. If the parameters *q*, *e* belong to the domains ***D*_1_** or ***D*_3_**, then, for any initial values (*R*, *P*), the system collapses (*R* → 0, *P* → 0);
2. If the parameters *q*, *e* belong to the domain ***D*_2_**, then, there exist domains of initial values in which the system tends to the parasite-free equilibrium *O*_2_ whereas, at other initial values, it tends to the trivial equilibrium *O* (collapse).

Therefore, the original model [26] has no non-trivial (*R* > 0, *P* > 0) equilibria whereby the replicator and the parasite could coexist and coevolve. A stable parasite-free equilibrium can exist only if 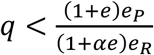 and necessarily exists if, additionally, *ae* < (*K* - 4*e*_*R*_)/(4*e*_*R*_)), that is, if the reproductive advantage of the parasite is low and the replicator possesses defense mechanisms that are efficient enough and not too costly to overcome the intrinsic advantage of the parasite.

The implications of these findings are the following: emergence of a genetic parasite as a (slightly) degraded copy of the “naÏve” (defenseless) replicator (i.e. when eqs. (1) apply and *e* =0) leads to the collapse of the system towards the trivial equilibrium (*R* = 0, *P* = 0). If the replicator already has a sufficiently advanced and not excessively costly defense mechanism (or is able to evolve it before the population collapse), the system could be stable in the parasite-free state as long as the defense mechanism persists. If the defense degrades over time, which is likely to be the case because most if not all defense mechanisms incur a non-zero fitness cost [31-34], the system will become vulnerable again. Therefore, such a system is inherently unstable. If the early history of (pre-)life included a primitive replicator stage, as the RNA World concept implies [17, 35, 36], it would be vulnerable to parasite-driven collapse, could not have persisted for a long time and necessarily would evolve into a different mode of replicator-parasite relationships that is not subject to the limitations imposed by eq. (1).

### Genetic parasite with an additional interaction with the replicator

Let us consider the model that differs from the model (1) by an additional effect of the replicator-parasite interaction on the replicator dynamics:

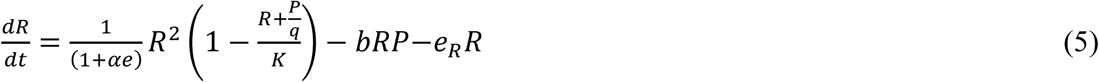

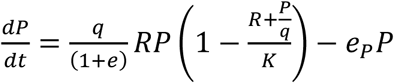

Such an effect could be interpreted as an extra penalty on the replication rate:

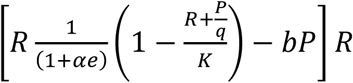

and/or as extra replicator degradation rate:

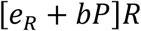

due to the replicator-parasite interaction.

The trivial equilibrium *O* (*R* = 0, *P* = 0), and the semi-trivial equilibria *O*_1_ and *O*_2_ (*R* > 0, *P* = 0), identified for the model (1) still exist in the system (5) under the same conditions. In addition, however, two new equilibria also can exist in the system, namely, *A*_1_(*R*^+^, *P*^*^) and *A*_2_(*R*^-^, *P*^*^) where

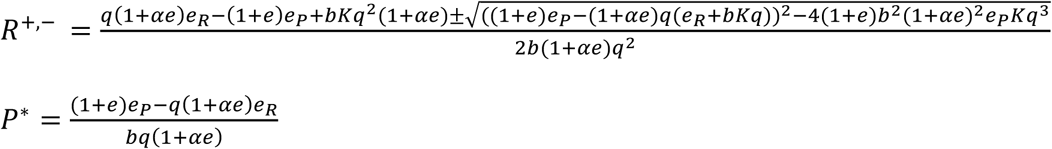

These equilibria are non-trivial (i.e. imply *P* > 0) only if 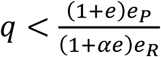; if 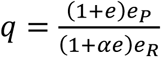, the equilibrium points *A*_1_ and *A*_2_ merge with the points *O*_1_ and *O*_2_ of eq. (3). Stability analysis, however, shows that, when these equilibria exist, both are unstable: *A*_1_ is an unstable node and *A*_2_ is a saddle (Figure 4a, b; see Additional File 2 for details). Therefore, although the inclusion of this extra interaction, corresponding to a more aggressive parasite compared to model (1), into the model does lead to non-trivial equilibria (i.e. replicator-parasite coexistence), they are unlikely to persist for on the evolutionarily relevant time scale.

**Figure 3.**
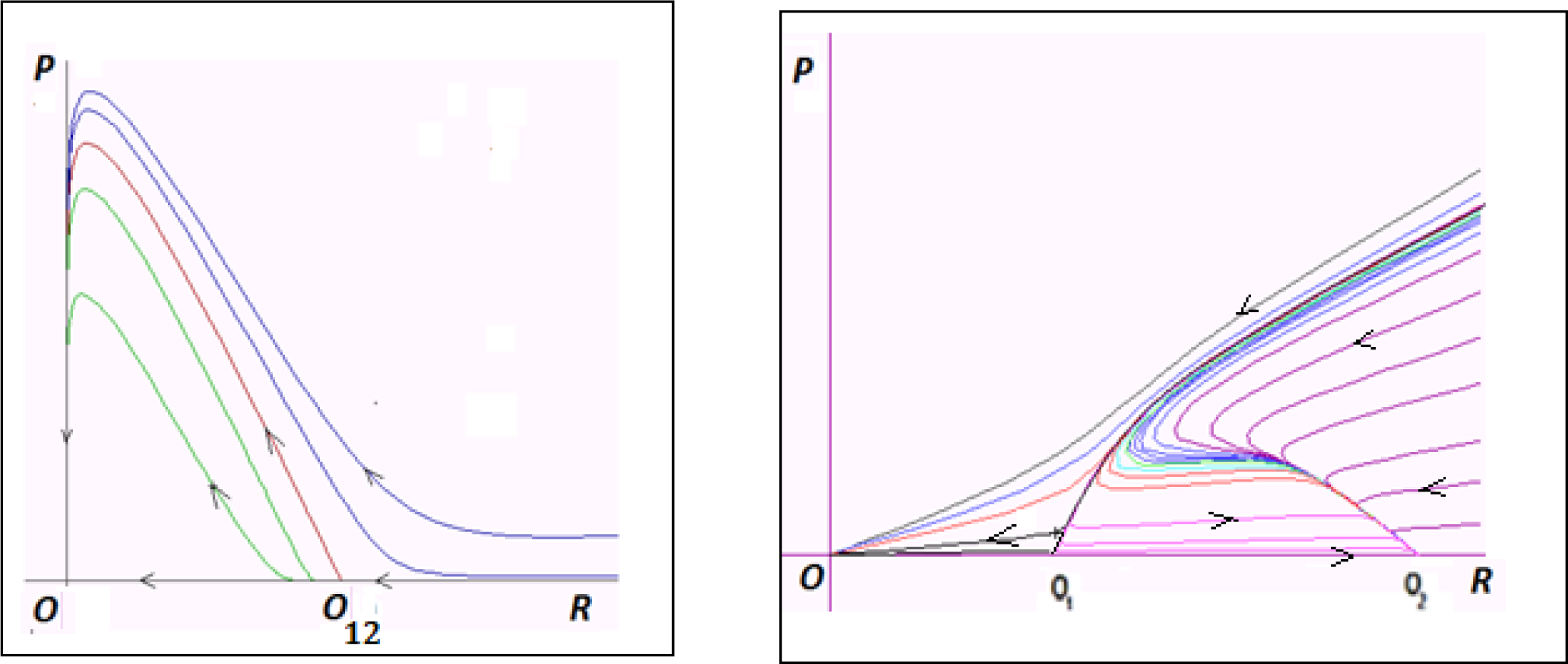
Phase portraits of the model (1) on the boundaries *B* (left panel) and *Q* (right panel) of the parameter-phase portrait.

**Figure 4.**
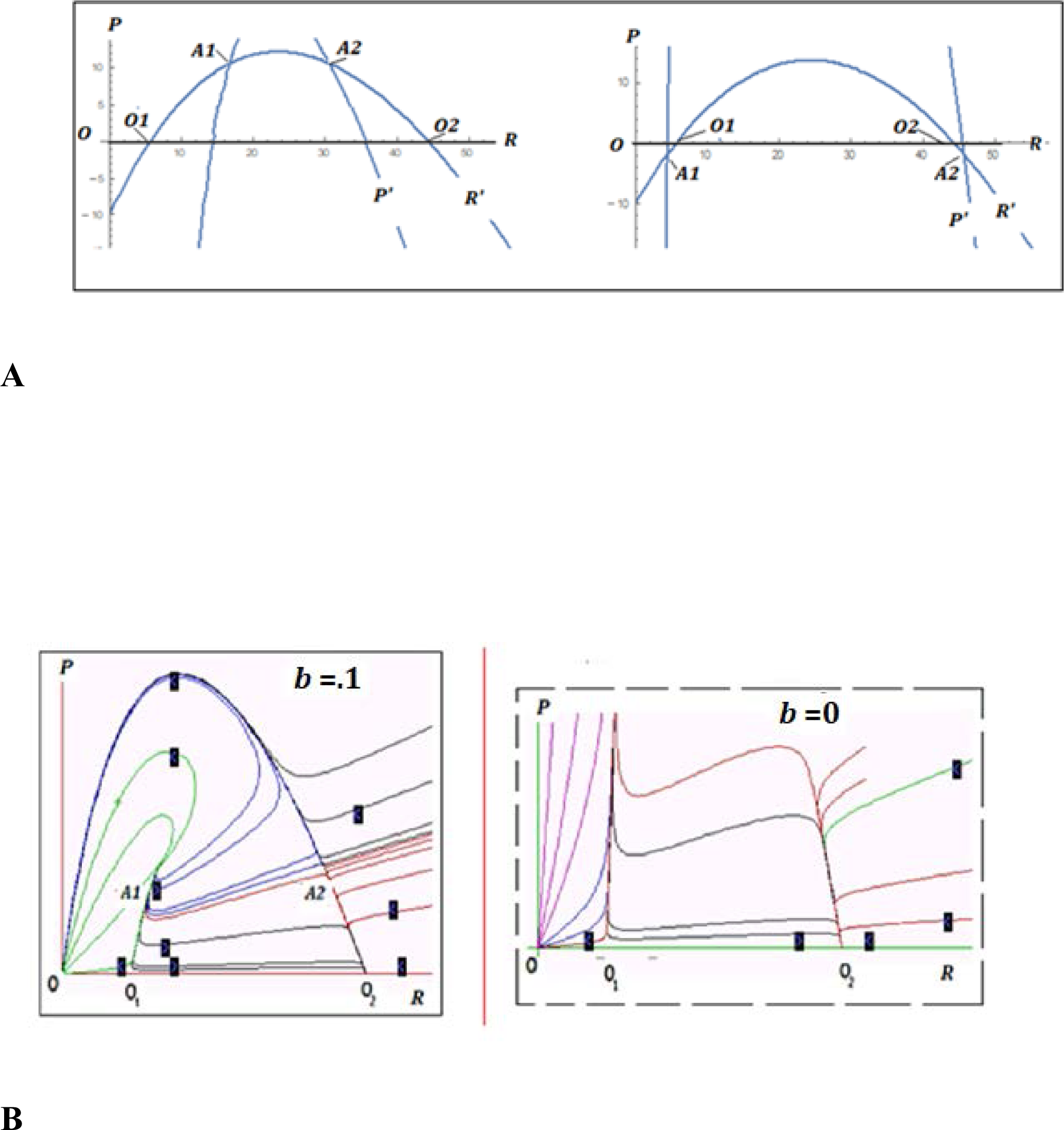
Characteristics of model (5). **A**. Null isoclines and equilibria. Left panel: positive non-trivial equilibria *A*_1_, *A*_2_, *q* = 4; right panel: no positive non-trivial equilibria, *q* = 10. **B**. Phase portraits. Left panel: phase portrait of the model (5; b>0) with unstable non-trivial equilibria *A*_1_ and *A*_2_. Right panel: phase portrait of the model (5; b=0, equivalent to model (1)), no non-trivial equilibria.

### A highly derived genetic parasite

Analysis of the model (1) shows that the equilibrium between the parasite and the replicator is unstable and largely hinges on the relative growth advantage of the parasite *q* and the efficiency of the host replicator’s defense mechanisms 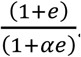. A parasite that has a greater advantage over the host replicator than the defense mechanisms can handle, overwhelms the system and drives it to collapse, whereas a less efficient parasite is eliminated by the host. Simply shrinking the parasite genome and hence increasing its replicative advantage (*q ≫* 1), while concomitantly reducing the impact on the replicator dynamics according to the expression 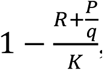, does not represent a viable path for the parasite towards the stable coexistence because the increase in the parasite replication efficiency overwhelms the attenuation of its deleterious effect. Moreover, as the analysis of the model (5) indicates, another intuitively plausible path to coexistence, through additional suppression of the replicator (allowing the parasite to escape elimination in the presence of the efficient defense mechanism, for example, by evolving antidefense mechanisms) does not work either. Although non-trivial equilibria can exist under this model, they are unstable.

More generally, the equilibria appear in the system as intersections of the isoclines (see eq. (2)). Thus, to ensure stable coexistence of the host and the parasite, the equations of the replicator and parasite dynamics should substantially differ from each other unlike those in model (1) (see Additional File 1). From the biological perspective, this means that the parasite cannot be, simply, a slightly modified variant of the replicator. Rather, the effects of the replicator and the parasite on each other’s replication should be substantially asymmetric and/or their interactions with the environment should be substantially different. One such possible modification is represented in the following model:

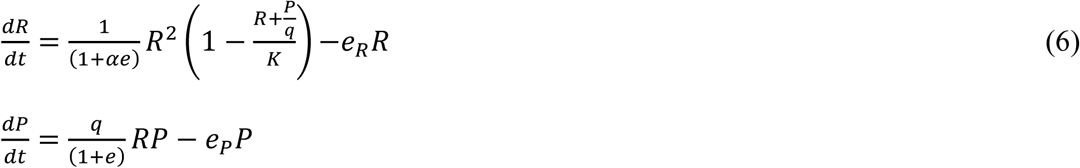

In the model (6), the parasite dynamics does not depend on the carrying capacity of the environment, whereas the consumption of the resources by the parasite continues to be a factor in the replicator dynamics 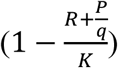. A biological model for this behavior is a highly specialized parasite that obtains the necessary resources from the host (effectively, for free) rather than directly from the environment. If the parasite is individually small enough compared to the host, as is the case for many viruses and transposons, at least those that parasitize on eukaryotes, its growth is not limited by the external environmental resources. The mere availability of the host ensures that the resources are sufficiently abundant and is the only limiting factor for the parasite reproduction.

Analysis of the model equilibria (see Additional File 3 for details) defines the following family of null isoclines for *P* and *R*:

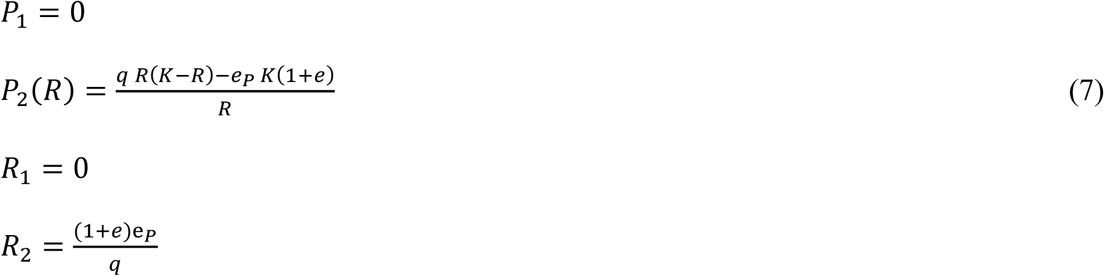

(Figure 5). The trivial equilibrium *O* (*R* = 0, *P* = 0), and the semi-trivial equilibria *O*_1_ and *O*_2_ (*R* > 0, *P* = 0), found in model (1) and defined by eq. (3), also exist in model (6) subject to the conditions defined by eq. (4). In addition, an equilibrium

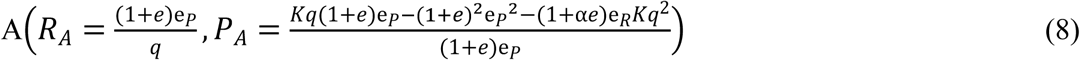

also exists. Under the condition 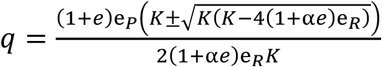, the equilibrium *A* is semi-trivial (*R* > 0, *P* = 0) and coincides with either *O*_1_ or *O*_2_. For the values of *q* defined by

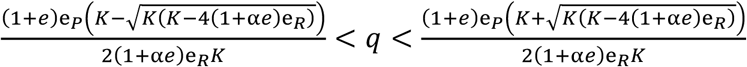

the equilibrium *A* is non-trivial (*R* > 0, *P* > 0).

**Figure 5.**
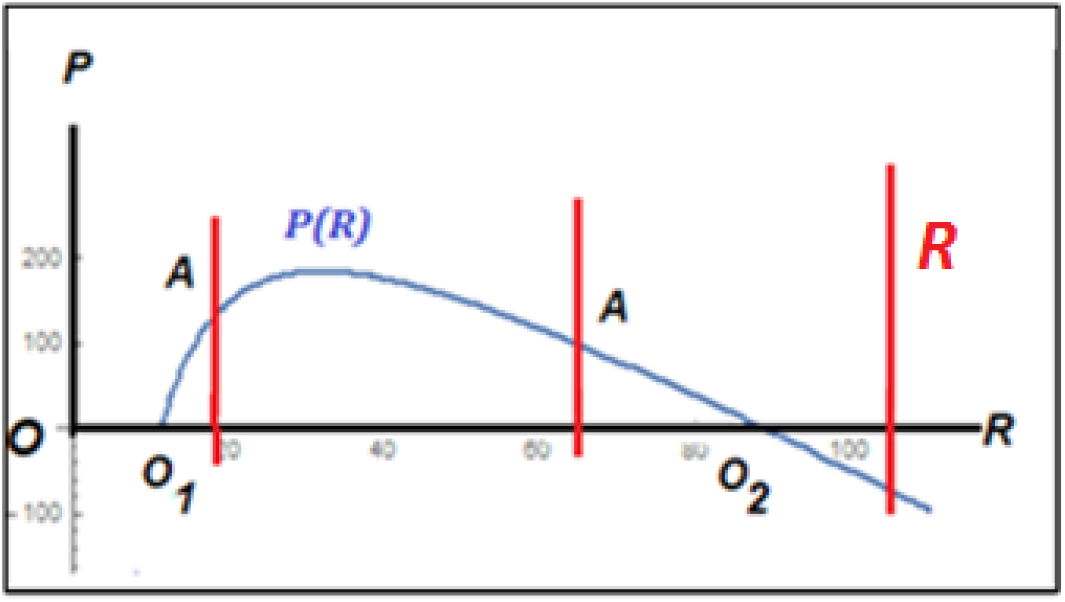
Null isoclines (eq. (7)) for model (6).

Analysis of the model (6) (see Additional File 3 for details) allows us to construct the parameter-phase portrait of the model (Figure 6a,b). Let us consider *q* and *e* as parameters of the model (6) whereas *K*, *α*, *e*_*R*_, *e*_*P*_ are arbitrary values of fixed coefficients. The portrait contains domains ***D*_1_**, ***D*_2_** and ***D*_3_** similar to those in model (1) (Fig.2), and 3 additional domains, ***D*_4_**, ***D*_5_** and ***D*_6_**.

**Figure 6.**
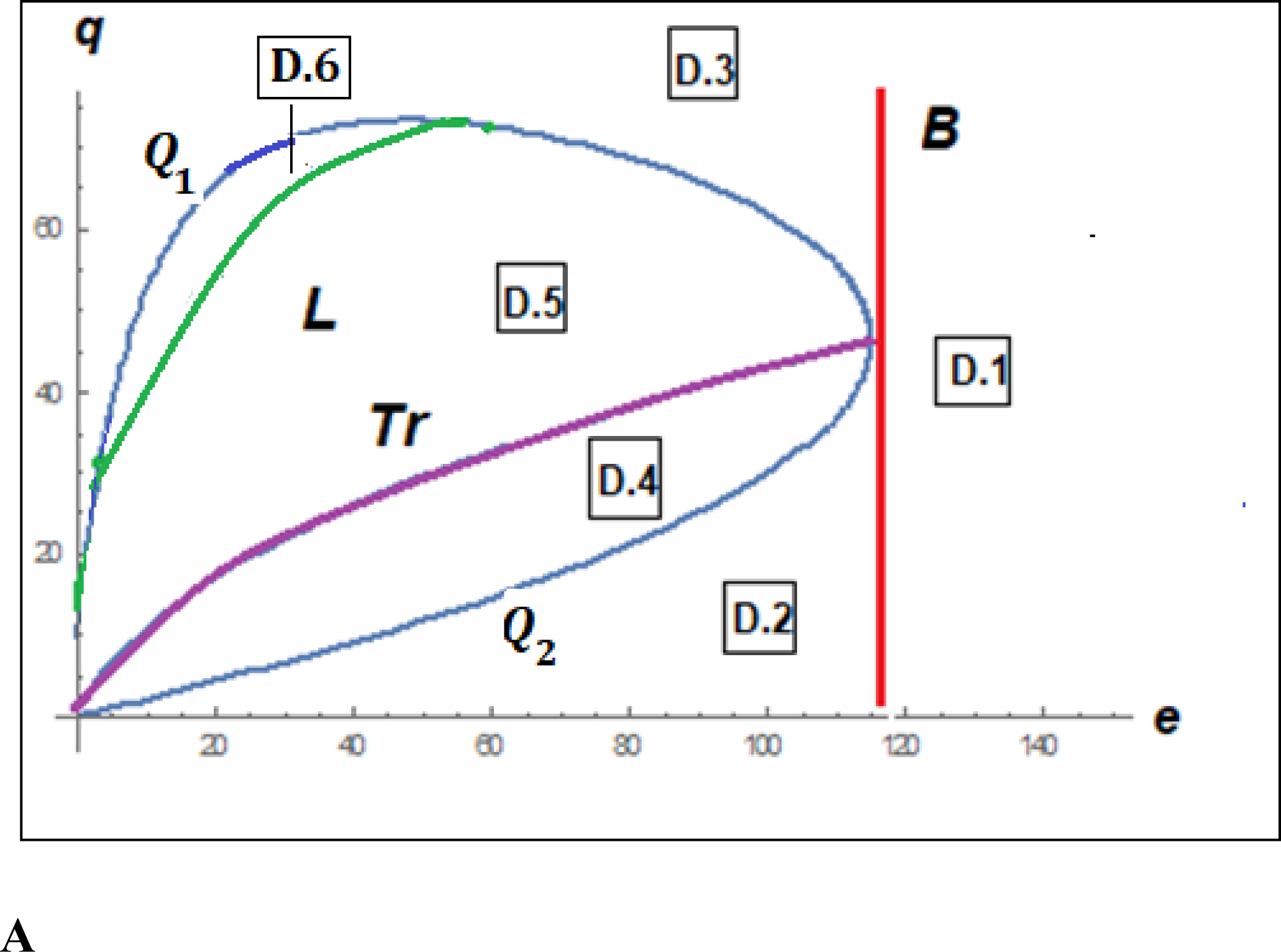
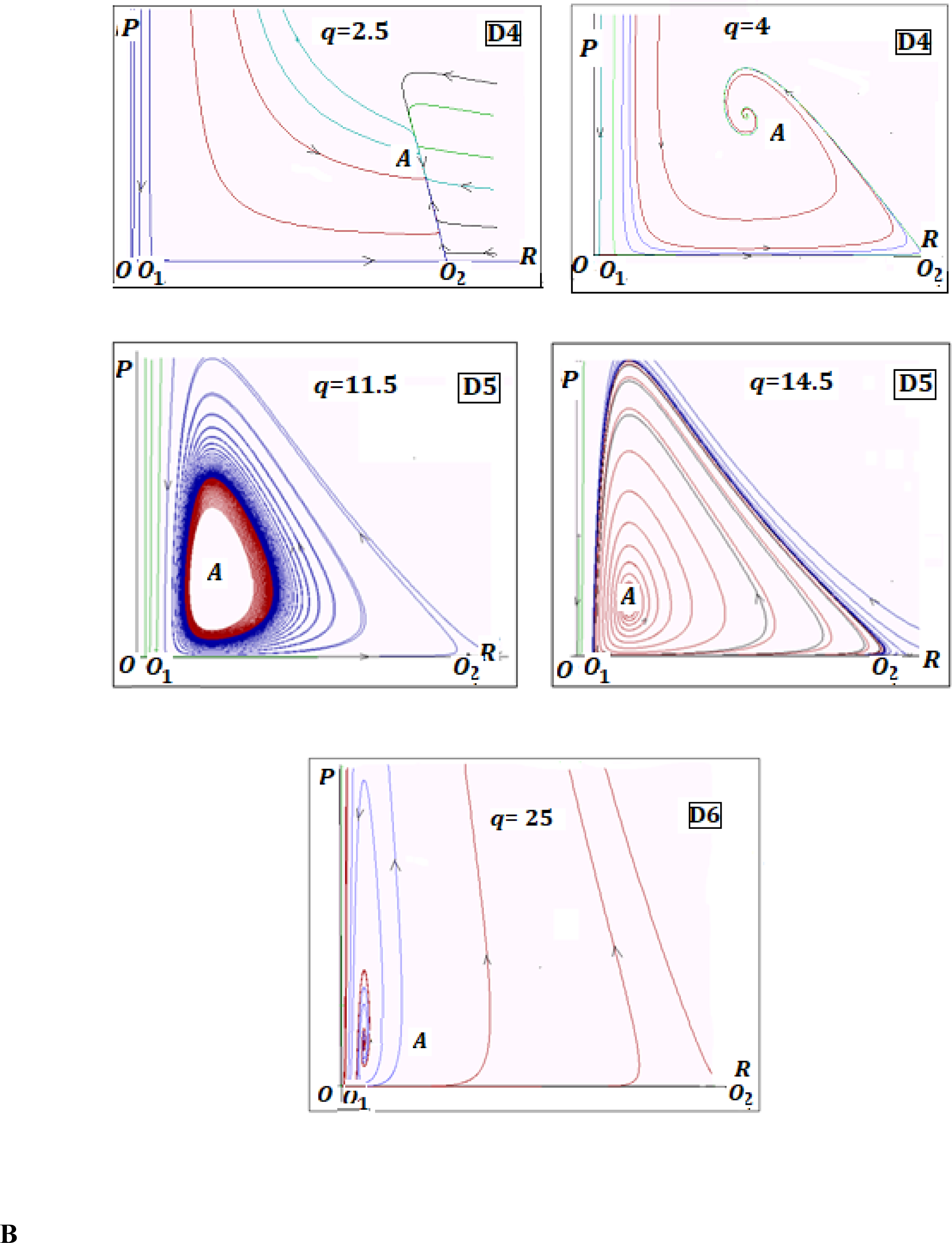
Characteristics of model (6). **A**. Parameter-phase portrait of model (6). ***D*_1_** – ***D*_6_** indicate domains with qualitatively different model behaviors. **B**. Examples of phase portraits of model (6) with a stable non-trivial equilibrium *A* (domain ***D*_4_**) and stable oscillations (domain ***D*_5_**); no non-trivial stable regimes in domain ***D*_6_**.

**Figure 7.**
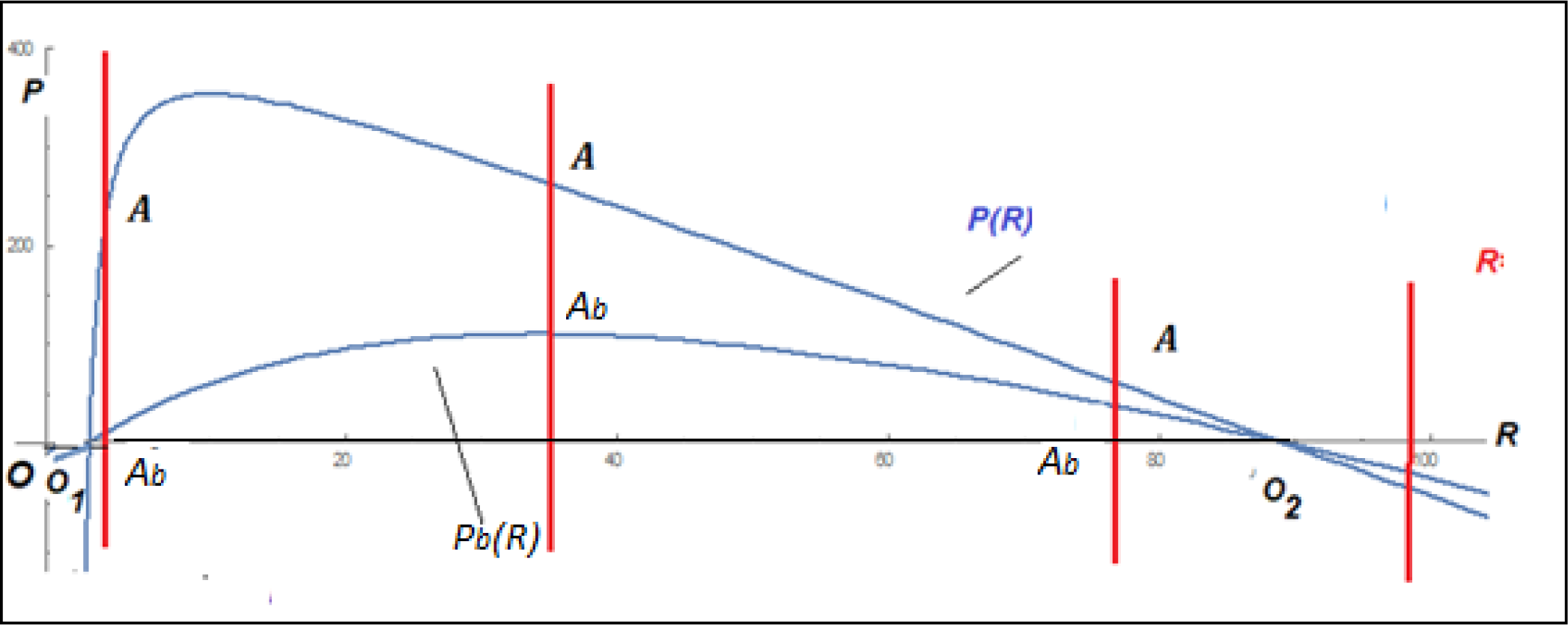
Null isoclines for Volterra-type models. The isocline *R*(*R*) corresponds to the model A3.1 which is the same as model (6);.the isocline *P*_*b*_(*R*) corresponds to the model A4.1 with b>0.

The boundaries between the parametric domains of model (6) are determined by the curves ***B***, ***Q*_1_**, ***Q*_2_**, ***Tr***, ***L*,** described in Additional File 3.

The domains have the following properties:

***D*_1_** (*e* > ***B***): only the trivial equilibrium *O* exists and is stable (in ***D*_1_**, the effective cost of the defense mechanisms *αe* is so high that the replicator population collapses even in the absence of the parasite);

***D*_*2*_** (*e* < ***B***, *q* < ***Q*_1_**): the trivial equilibrium *O* is stable; the semi-trivial equilibrium *O*_1_ is unstable; the semi-trivial equilibrium *O*_2_ is stable; no other equilibria exist (in ***D*_2_**, the efficiency of the defense mechanisms is sufficient to eliminate the parasite as long as the population of the replicator itself is sustainable);

***D*_3_** (*e* < ***B***, *q* > ***Q*_2_**): the trivial equilibrium *O* is stable; both semi-trivial equilibria *O*_1_ and *O*_2_ are unstable; no other equilibria exist (in ***D*_3_**, the parasite overwhelms the defense mechanisms of the replicator and thus collapses the system);

***D*_4_** (*e* < ***B***, ***Q*_1_** < *q* < ***Tr***): the trivial equilibrium *O* is stable; both semi-trivial equilibria *O*_1_ and *O*_2_ are unstable; the non-trivial equilibrium *A* is a stable node or focus (in ***D*_4_**, the parasite and the replicator can coexist at a stable equilibrium, Figure 6b);

***D*_5_** (*e* < ***B***, ***Tr*** < *q* < min(***L*, *Q*_2_**)): the trivial equilibrium *O* is stable; both semi-trivial equilibria *O*_1_ and *O*_2_ are unstable; the non-trivial equilibrium *A* is unstable but is surrounded by a stable limit cycle (in ***D*_5_**, the parasite and the replicator can coexist in a stable oscillation regime);

***D*_6_** (*e* < ***B*, *L*** < *q* < ***Q*_2_**): the trivial equilibrium *O* is stable; both semi-trivial equilibria *O*_1_ and *O*_2_ and the non-trivial equilibrium *A* are unstable (in ***D*_6_**, the defense mechanisms of the replicator are not sufficient to keep the parasite in check; although other equilibria exist, they are unstable, so that the system would collapse upon a perturbation).

Overall, the system collapses in domains D1, D3, and D6; the replicators and parasites can coexist in domains D4 and D5; and, the system is bistable in domains D2, D4 and D5, that is, its final behavior critically depends on the initial values of *P* and *R*.

Thus, under the model (6), there exists an intermediate regime in which the parasite efficiency is roughly balanced by the efficiency of the replicator defense mechanisms under which the replicator and the parasite can coexist indefinitely in a stable equilibrium or a limit cycle. Unlike in the model (1) that lacks such a regime, the parasite in the model (6) is a highly derived state. In such a state, the parasite replication and survival directly depend only on the host (replicator) availability but not on the environment (the parasite equation in (6) lacks the 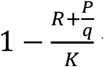 term). Models with the parasite being closely similar to the replicator in its interaction with the environment (that is, with the parasite replication equation closely resembling that of the replicator), such as the model (1), cannot support stable non-trivial equilibria (compare the families of null isoclines in eqs. (2) and (7)). A variant of the model (6) with an additional effect of the replicator-parasite interaction on the replicator dynamics shows qualitatively the same behavior as model (6) (see Additional File 4).

### Concluding remarks

We show here that, in order to produce stable equilibria, models of host-parasite interaction cannot be too simple. The parasite must not be too closely similar to the host in its reproduction strategy. More specifically, stable coevolution becomes possible in models where only the reproduction of the host but not that of the parasite depends on the carrying capacity of the environment. From a biological perspective, a successful parasite has to rely on the host not only for replication but also for building blocks and energy. Perhaps, these results go some way to explain why, to the best of our current knowledge, no genetic parasites have ever captured neither full-fledged biosynthetic pathways, including the translation system, nor the molecular machinery for energy production.

Although the analyzed models are too simple to be of much direct relevance to extant cases of host-parasite coevolution, they are likely to be relevant for early stages of (pre)life evolution, within the RNA World, the leading current scenario for the origin of life [37-39], and, at the subsequent stages, when DNA and proteins came to the scene, and different replication strategies evolved. Importantly, the replicators in this model are not abstract information carriers but self-sustaining reproducers that extract energy and building blocks from the environment, supporting both the host and the parasite [17, 35].

The results of this work imply that primordial replicators have made innumerable “false starts” whereby host-parasite systems collapsed under the unchecked parasite pressure. Only in more evolved systems, where the capacity of the parasite to outcompete the host is balanced by defense mechanisms and, conversely, the ability of the host to eliminate the parasites is undermined by the cost of defense, stable coevolution became possible. Such coevolution between hosts and parasites is likely to be an essential driver of the evolution of biological complexity, and more specifically, of major transitions in evolution [15]. Hence any biological systems, in which a stable host-parasite coevolution regime failed to evolve, would be evolutionary dead ends.

We considered here only homogenous, well-mixed host-parasite systems. In computer simulations, compartmentalization has been shown to lead to stable co-evolutionary regimes. Most likely, compartmentalization of replicator ensembles had been part and parcel of the evolution of life from its earliest stages on, and could be considered a form of host defense, perhaps, the simplest one [18, 19]. Clearly, compartmentalization is a major path to the emergence of diversity and complexity. Further development of the models described here, in particular, by explicitly allowing evolution of the parameters defining the behavior of the parasites and replicators, will be of interest.

## Declarations

### Ethical Approval and Consent to participate

Not applicable

### Consent for publication

Not applicable

### Availability of supporting data

Supporting data are available as additional files.

### Authors’ contributions

FB and GPK performed research; FB, GPK, YIW, MIK, and EVK analyzed the results; GPK, YIW, and EVK wrote the manuscript that was edited and approved by all authors.

### Authors’ information

Faina Berezovksaya is at the Department of Mathematics, Howard University, Washington, DC 20059, USA; Georgy P. Karev, Yuri I. Wolf, and Eugene V. Koonin are at the National Center for Biotechnology Information, National Library of Medicine, National Institutes of Health, Bethesda, MD 20894, USA; Mikhail I. Katsnelson is at the Institute for Molecules and Materials, Radboud University, 6525AJ, Nijmegen, Netherlands.

## Additional files

Additional File 1

Mathematical Appendix 1

Additional File 2

Mathematical Appendix 2

Additional File 3

Mathematical Appendix 3

Additional File 4

Mathematical Appendix 4

